# Long-read sequencing reveals novel structural variation markers for key agronomic and quality traits of soybean

**DOI:** 10.1101/2024.01.09.574864

**Authors:** Zhibo Wang, Kassaye Belay, Joe Paterson, Patrick Bewick, William Singer, Qijian Song, Bo Zhang, Song Li

## Abstract

In plant genomic research, long read sequencing has been widely used to detect structure variations that are not captured by short read sequencing. In this letter, we described an analysis of whole genome re-sequencing of 29 soybean varieties using nanopore long-read sequencing. The compiled germplasm reflects diverse applications, including livestock feeding, soy milk and tofu production, as well as consumption of natto, sprouts, and vegetable soybeans (edamame). We have identified 365,497 structural variations in these newly re-sequenced genomes and found that the newly identified structural variations are associated with important agronomic traits. These traits include seed weight, flowering time, plant height, oleic acid content, methionine content, and trypsin inhibitor content, all of which significantly impact soybean production and quality. Experimental validation supports the roles of predicted candidate genes and structural variant in these biological processes. Our research provides a new source for rapid marker discovery in crop genomes using structural variation and whole genome sequencing.

## Main

Decades of research have shown that structural variations (SVs) including deletions, insertions, duplications, and chromosomal rearrangements, are important in plant evolution and agriculture, affecting traits such as shoot architecture, flowering time, fruit size, and stress resistance ^1,2^. Soybean is an important source of food, protein, and oil, its whole genome sequence is critical for the improvement of these traits in soybean breeding and research programs, as it allows the discovery of genes and their functions, as well as the development of genetic markers for selection. The wm82v2 assembly was built primarily on short reads and Sanger sequence, the Wm82v2 assembly was primarily Sanger-based, and gap-filling in v3 and v4 used PacBio-based BAC assemblies targeted to the gap regions. At least in the V4, long-reads have also been in cooperated ^3^. The third generation long-read sequencing technology has been widely adopted in the research community, since it can facilitate the identification of SVs that are unable to be captured by short-read sequencing technology ^4-6^.

Our objective was to assess the presence, distribution, and the phenotypic effects of large-scale structural variants (> 50bp indels) ^7,8^ in soybean genotypes. To achieve this goal, we applied Oxford Nanopore Long-Read Sequencing Technology (ONT) to sequence 28 representative genotypes of both grain-type soybean and vegetable soybean (edamame) as well as the reference variety, William 82 (W82, Table S1). This germplasm compilation includes grain-type soybeans such as Hutcheson and V12-4590 used in livestock feeding, food-grade soybeans like zizuka (natto), V10-3653 (soy milk and tofu producing variety), MFS-561 (sprout variety), and vegetable soybeans (edamame) including VT-Sweet and V16-0565 ^9-12^. These genotypes feature diverse traits such as high methionine content, high yield, varying seed protein levels, distinct plant heights (both high and low), and high oleic acid content, and different seed weights (Figure 1 and Table S1).Our study doubled the number of soybean genotypes that were re-sequenced by a long-reading sequencing approach ^4^. For the 29 soybean genotypes, we collected a total of 5.6 Tb of long-read raw sequencing data with an average of 30X genome sequence coverage and an average read length N50 of 11.1 kbp. The recently published Wm82.a4.v1 soybean reference genome ^3^ was utilized to align raw sequencing reads using minimap2 ^13^. The resulting aligned reads were sorted and indexed using samtools ^14^ and structural variants were called with Sniffles ^15^. Called structural variants were then filtered (defined in this study as >50 bp and < 15,000 bp) and all SV calls for all 29 genotypes were merged. A total of 365,497 SVs were identified and has been uploaded for widespread use within the soybean community as a supplementary dataset.

**Figure 1.**
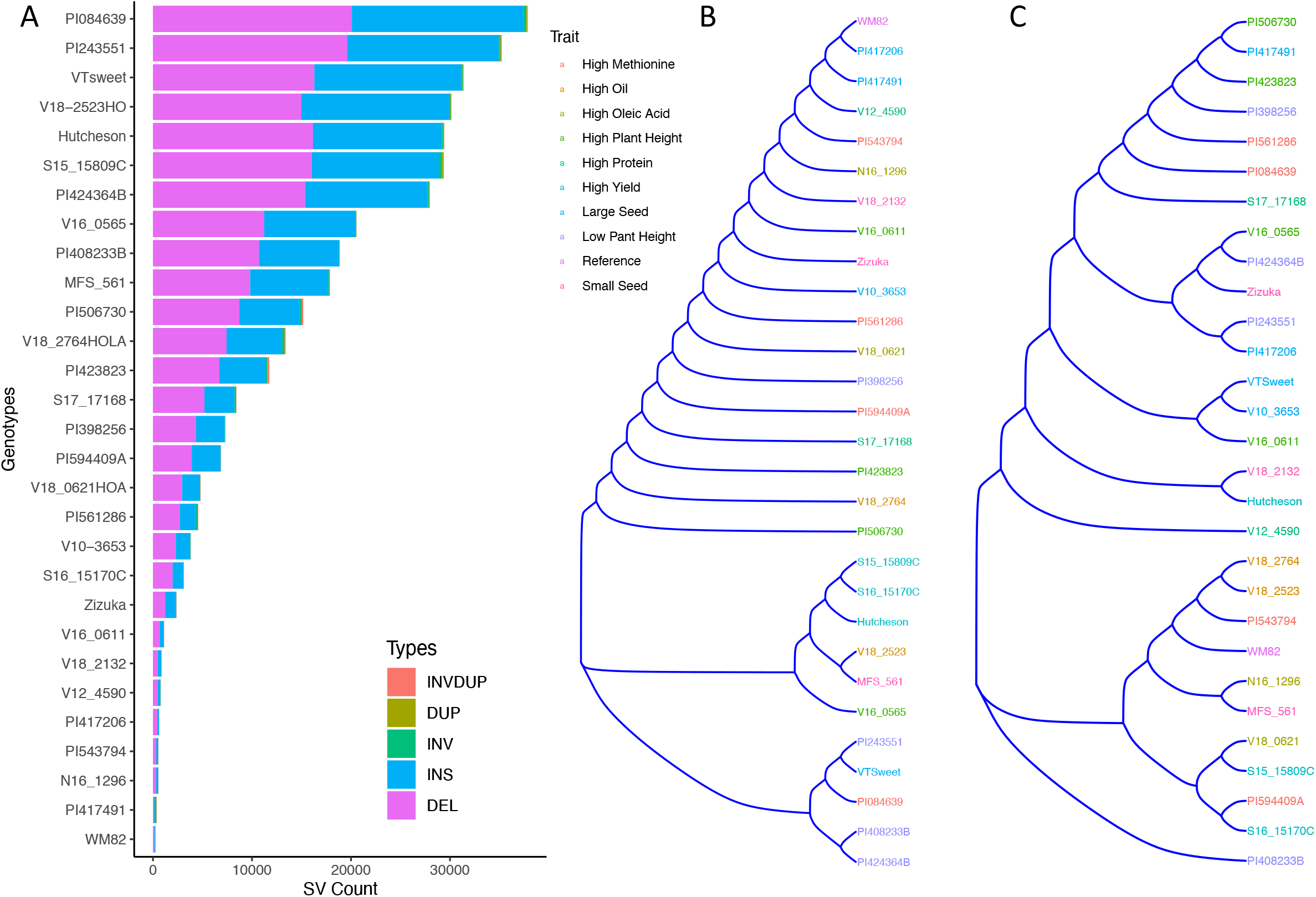
The clustering of 29 soybean/edamame genotypes based on their structural variant presence/absence matrix. (A) Stacked bar graph showed structural variant numbers and types from 29 soybean genotypes. (B) Hierarchical clustering dendrogram of structural variant presence/absence matrix across 29 soybean genotypes, with colors corresponding to phenotypic traits.

These 29 soybean genotypes had SVs between 253 (W82) and 37,863 (PI084639). PI084639, PI243551, V18-2523HO, and VT Sweet, carried the most structural variation relative to the WM82 reference genome (Table S1). Insertions and deletions were the most common SV types in all genotypes, accounting for more than 95% of the SVs (Table S1 and Figure 1A). We also found hundreds of inversions, duplications, and translocations among the accessions (Table S1 and Figure 1A). Cluster analysis based on the SV presence/absence matrix showed that the genotypes were clustered into three major groups (Figure 1B). Interestingly, genotypes sharing similar representative traits did not form clusters, except for the notable observation that the genotypes we specifically chose (Hutcheson, S16-15170C, and S15-15809C) for high yield were found to cluster together (Figure 1B). The two accessions selected as the representatives of low plant height were clustered together but PI424364B had a much greater number of SVs as compared to PI408233B (Table S1, Figure 1A and 1B). In addition, the same 29 varieties were genotyped using 6K beadchip assay with 6K SNPs to generate a SNP-based phylogenetic tree (Figure 1C), which showed a different relatedness as compared to the tree based on SV presence/absence. This result suggests that genetic relatedness of soybean varieties based on structural variation represents a significant deviation from that deduced from SNPs.

We further evaluated SVs’ distributions based on their length and location. The length distribution of SVs showed that majority of SVs (61.3%) had the length ranging at 501-1500 bp, followed by 17.6%, 9.9 %, 8.0 %, 3.2% of SVs within 50-100 bp length, 101 to 200 bp, 201-500 bp, and above 1501 bp, respectively, (Figure 2A, Table S2). Compared to the SNPs, SVs can cause large-scale perturbations of cis-regulatory regions and are therefore more likely to quantitatively change gene expression and alter related agronomic phenotypes ^1,16^. For instance, a tandem triplication over the AMTE1 genes is reported to be associated with aluminum resistance in maize ^17^ and an insertion of Ty1/copia-like retrotransposon disrupted the expression level of E4 and caused insensitivity under long day conditions in soybean ^18^. In our analysis, we found that 77% of the total SVs (285, 760 structural variants) are located within intergenic 3-15 kb region (Figure 2B and Table S3). To confirm whether our candidate SVs affected gene expression, and more importantly, on plant phenotypes, we performed experimental validations and statistical analyses to verify the functions of the identified SVs.

**Figure 2.**
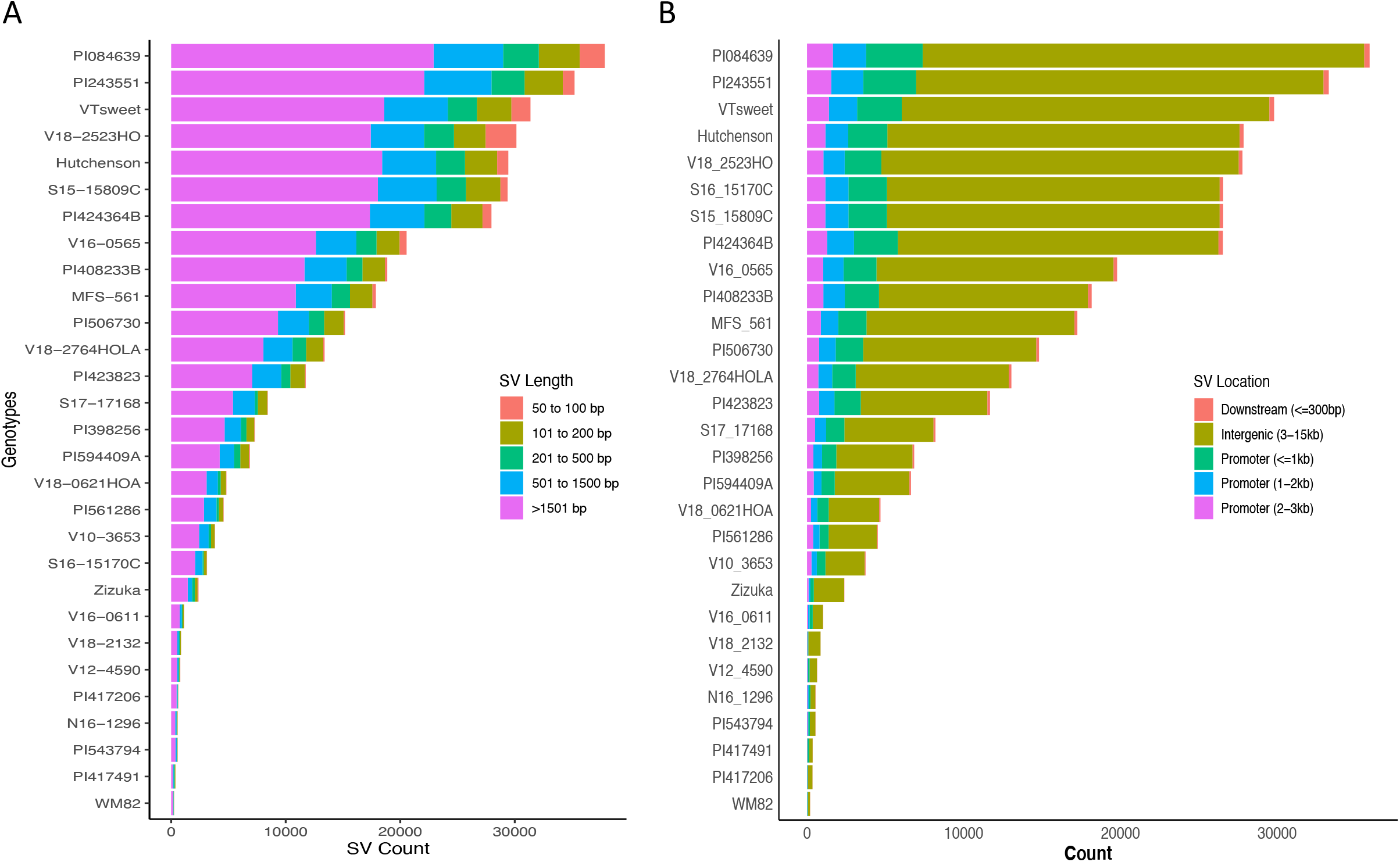
The distribution of SVs across 29 soybean genotypes. (A) Stacked bar graph showed structural variant length distributions, (B) Stacked bar plot for structural variant distributions across different genomic regions.

In light of previous findings associating QTLs, SNPs, and gene models with seed quality and agronomic performance in soybeans, our study focused on SVs derived from the current research. We specifically selected an SV located within a QTL region known to impact soybean seed Kunitz trypsin inhibitor (KTI) for functional validation ^19^. Soybean meal provides an excellent source of protein in animal feed since it is rich in amino acids with a high nutritional profile. However, the digestibility of soy protein can be severely impacted by KTI, which can restrain the function of trypsin, a critical enzyme that breaks down proteins in the digestive tract^20^. Traditional heating processes used in soybean meal production deactivate KTI, yet this method not only reduces the meal’s nutritional value due to amino acid degradation but also escalates energy costs by 25%. Raising low-KTI or KTI-free soybeans on farms creates a unique market opportunity for integrated crop and livestock farmers, increasing their farm’s profitability. A major QTL at chromosome 8 in the soybean genome was reported to harbor 13 KTI homologue genes and associate with low KTI in the mapping population ^19^. The present study found a deletion of 1443 bp in the downstream sequence of two KTI genes (Gm08g342200 (KTI7), Gm08g342300 (KTI5)) in the two inbred lines, V12-4590 and S17-17168 (Figure 3A), from Virginia and Arkansas, respectively. The deleted SV was visualized by integrative genomics viewer (IGV) and PCR using the genomic DNAs as templates (Figure 3B and 3C). The real-time PCR showed that the expressions of the two genes were both reduced in the seeds of these two lines in contrast with those in WM82 (Figure 3D). The seed KTI content in the two lines was significantly lower than that in WM82 (Figure 3E). Here, V12-4590, also names by VT Barrack, was purposely bred as a low KTI soybean variety while S17-17168 has not been reported to be a low KTI variety. Our results revealed a new genetic variation regulating the KTI genes’ expressions in the soybean seeds.

**Figure 3.**
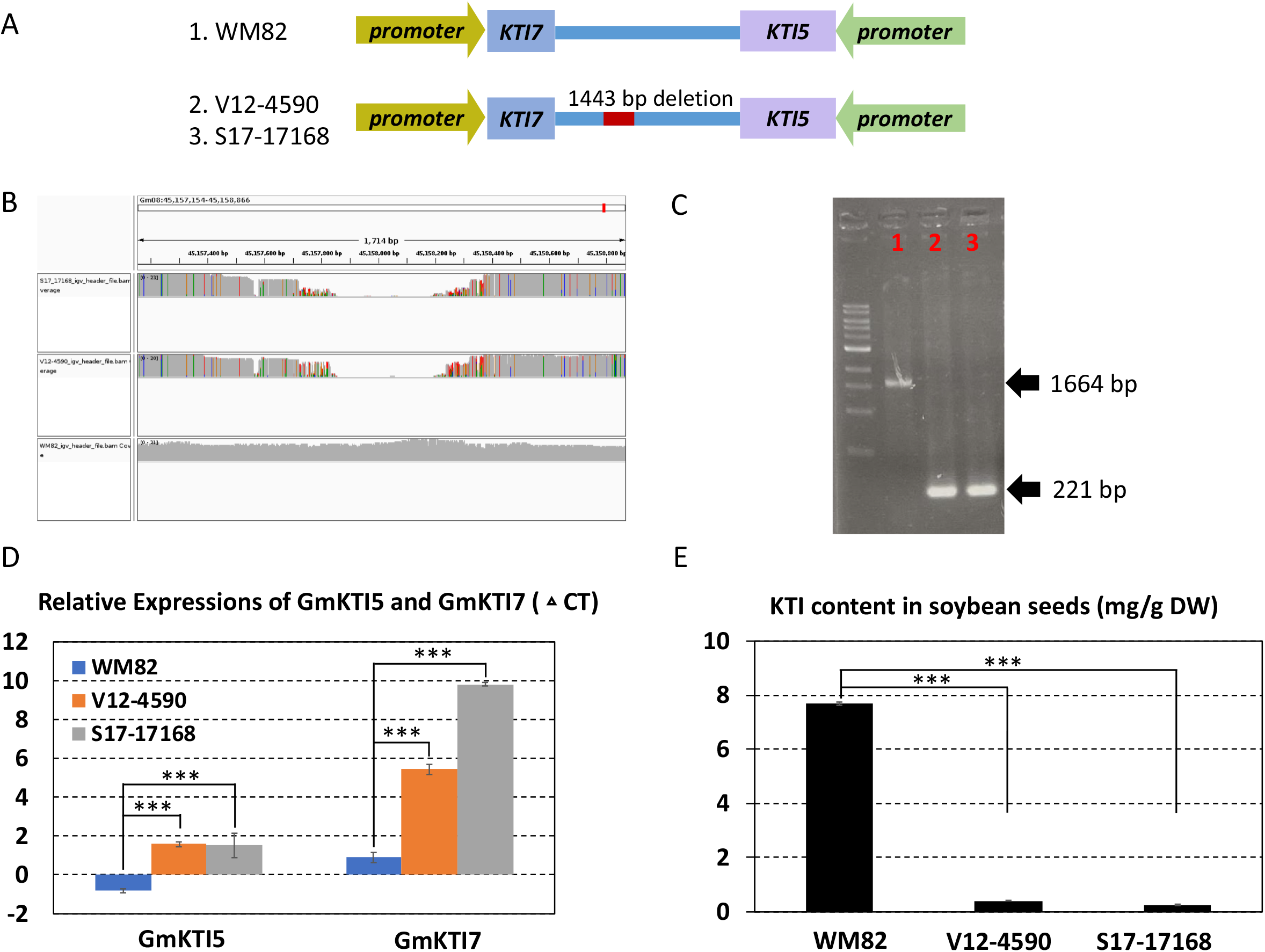
A structural deletion (1443 bp) was found to locate between two KTI genes (Gm08g342200 and Gm08g342300); validated to lower the expression levels of these two genes and reduce the KTI content in the soybean seeds. (A) The scheme displayed the location of this structural deletion. The orientations of two genes were opposite, and structural deletion was located at 3’ UTR of these two genes. (B) 1443 bp deletion was detected in two lines, V12-4590 and S17-17168 other than William 82 (WM82). (C) Agarose gel image of PCR products spanning the deletion polymorphism. M:1 kb ladder; 1: WM82; 2: V12-4590; 3: S17-17168. As expected, WM82 exhibits 1664 bp PCR product, whereas the other two genotypes all show a 221 bp amplicon. (D) Real-time PCR was utilized to evaluate the expression levels of Gm08g342200 (KTI 7) and Gm08g342300 (KTI 5) in the seeds of three genotypes. With the structural deletion, the expressions of these two genes in the seeds of V12-4590 and S17-17168 dramatically declined as compared with WM82. (E) The KTI contents in the seeds of WM82, V12-4590 and S17-17168 were assessed by HPLC. Consistent with the expression result, the KTI content in the seeds of V12-4590 and S17-17168 was dramatically less than that in the seed of WM82.

Despite significant advancements, several unexplored facets persist within soybean genetics, where structural variants (SVs) potentially wield considerable influence. In order to harness the potential of the SVs called by present study, we employed Chi-squared test followed by Benjamini-Hochberg adjusted p-value thresholds to pinpoint candidate genes associated with six key traits crucial in the soybean industry, including (1) protein content, (2) oleic acid content, (3) methionine content, (4) seed weight, (5) plant height, and (6) flower color (Table S4 and Table S5). From this pool of traits and genes, we chose two candidate gene models for further validation.

A substantial increase in seed size was a major feature of soybean domestication, especially for the vegetable soybean, edamame. According to the annotation in Soybase ^21^, the maturation-associated protein 1 (MAT1, Gm07g090400) gene was associated with seed size. Our results found a 319 bp deletion in the promoter region (−448 bp to TSS) of MAT1 in two food grade soybean varieties, V10-3653 (food grade soybean) and V16-0524 (VT sweet, the edamame variety) (Figure 4A). The statistical analysis displayed that the SV locating with MAT1 has a p-value of 1.26E-02, showing a significant association with seed size of soybean seeds (Table S4). The deleted SV was visualized by IGV and confirmed by PCR using the genomic DNAs as templates (Figure 4B and C). With the deletion, the expression of MAT1 gene in the seeds of these two varieties was much higher than that in WM82 (Figure 4D). Consistently, the seed weights of the two lines were also much higher than the WM82 seed (Figure 4E). The regulating molecular mechanism through the present deletion will be further investigated.

**Figure 4.**
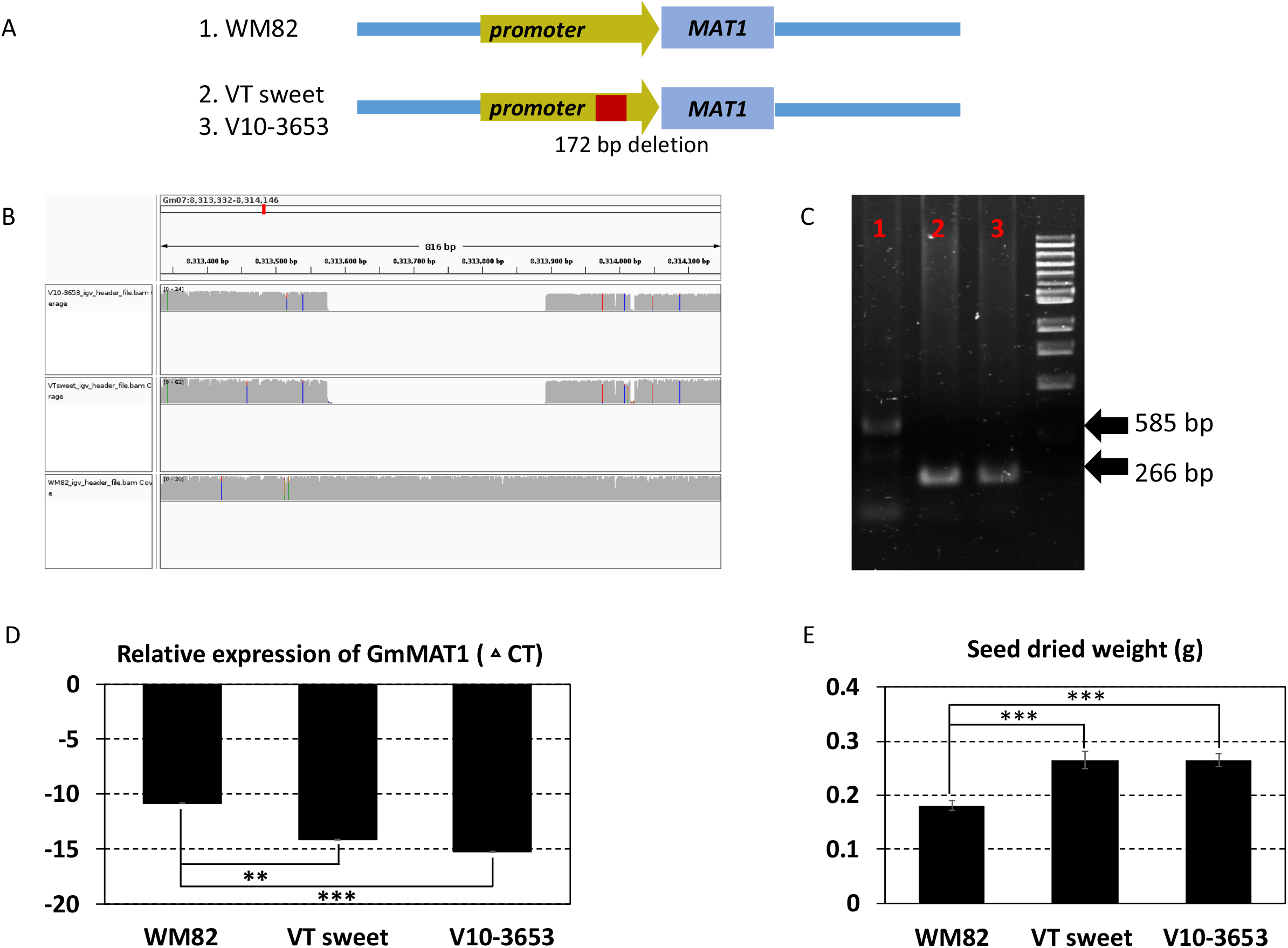
A structural deletion (319 bp) was found to locate at the promoter region of MAT1 (Gm07g090400); validated to enhance its expression level and the size of the soybean seeds. (A) The scheme displayed the location of this structural deletion. (B) 319 bp deletion was detected in two accessions, VT sweet and V10-3653, other than William 82 (WM82). (C) Agarose gel image of PCR products spanning the deletion polymorphism. M:1 kb ladder; 1: WM82; 2: VT sweet; 3: V10-3653. As expected, WM82 exhibits a 585 bp PCR product, whereas the other two genotypes all show a 266 bp amplicon. (D) Real-time PCR was utilized to evaluate the expression level of MAT1 in the seeds of three genotypes. With the structural deletion, the gene’s expression in the seeds of VT sweet and V10-3653 dramatically increased in comparison with WM82. (E) The seed size of WM82, V10-3653 and VT sweet were assessed by the seed weight, where the seed weight of WM82 was significantly lower than V10-3653 and VT sweet seeds.

The optimal height for current commercial soybean cultivar contributes to higher yields through improved resistance to lodging, with shorter or taller stands leading to yield reductions ^22^. Here, we found a deletion of 173 bp in the promoter region (−3688 bp to TSS) of EARLY FLOWERING 3 (ELF3, Gm04g050200) gene in three low plant height accessions including PI423464B, PI408233B, and PI243551 (Figure 5A). The statistical analysis showed that the SV locating at the promoter region of ELF3 has a p-value of 3.39E-02, suggesting a marginally significant effect on the plant height of soybean plants (Table S4). All the three accessions belong to the maturity group IV. The presence of the SV was confirmed by IGV and PCR using the genomic DNAs as templates (Figure 5B and 5C). ELF3 was reported as a core component of the circadian clock in the evening complex ^23^. Hence, we speculated the alteration on the expression of ELF3 might be responsible for the early flowering and relatively low plant height of these accessions. The regulating molecular mechanism through the deletion will be studied for comprehensive understanding of the functions of ELF3 in the circadian clock and soybean maturity.

**Figure 5.**
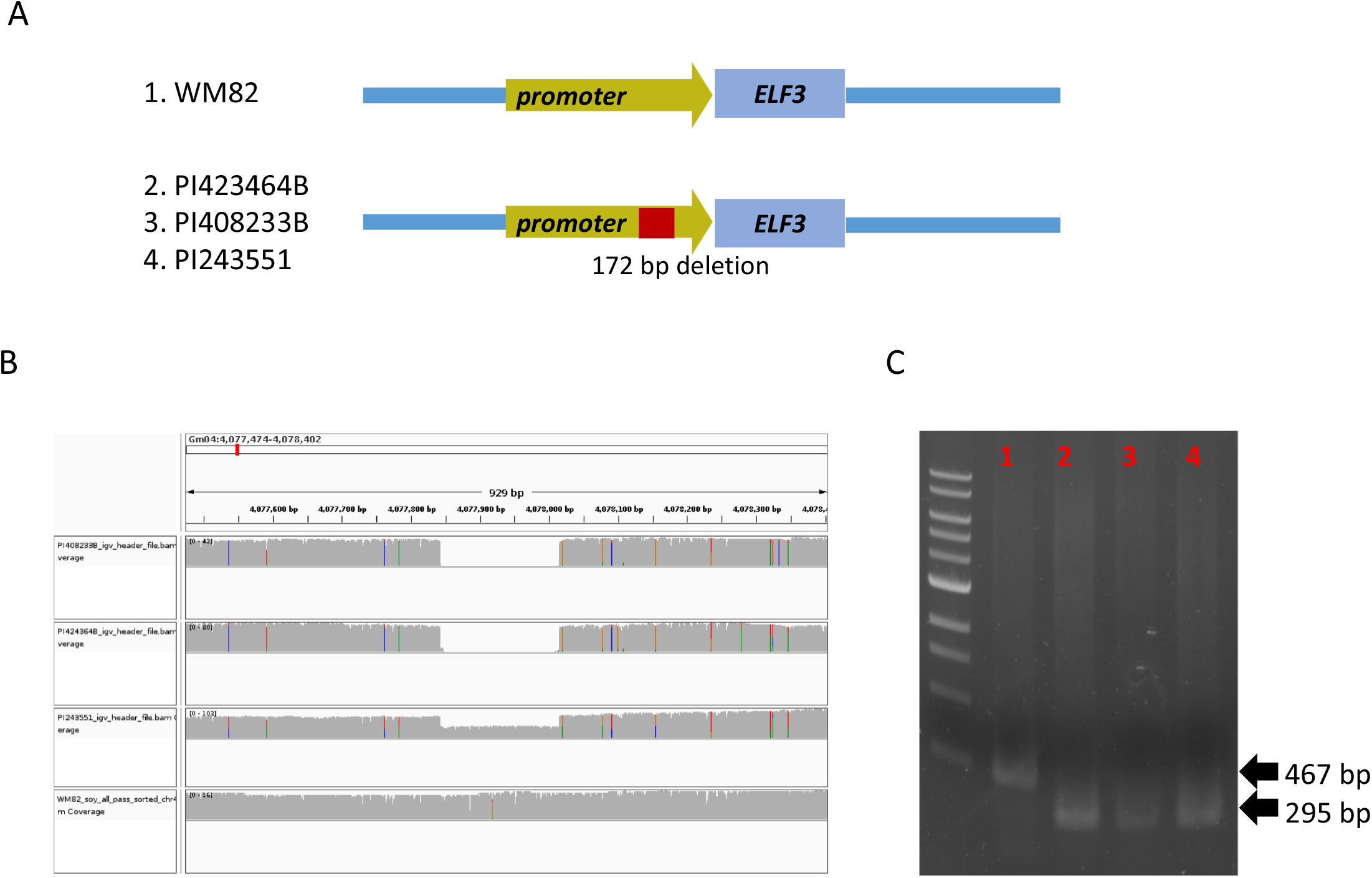
A structural deletion (172 bp) was found to locate at the promoter region of ELF3 (Gm04g050200), which might be associated with flowering time and maturity of soybean plants. (A) The scheme displayed the location of this structural deletion. (B) 172 bp deletion was detected in two accessions, PI423464B, PI408233B, PI243551, but not WM82. (C) Agarose gel image of PCR products spanning the deletion polymorphism. M:1 kb ladder; 1: WM82; 2: PI423464B; 3: PI408233B; 4: PI243551. As expected, WM82 exhibits a 467 bp PCR product, whereas the other three genotypes all show a 295 bp amplicon.

In conclusion, by integrating ONT to re-sequence 29 genotypes of grain type soybean and edamame, we identified novel structure variations that have the potential to reveal the basic genetic mechanism associating with important agronomic traits in soybean. It is fundamental to uncover the mechanism underlying these complex traits. These SVs can be also applied to molecular breeding via CRISPR/Cas9-mediated gene editing and to develop markers for soybean breeding selections. Our results provide a new pipeline for understanding basic genetics and rapid marker discovery in crop genomes using structural variation and whole genome sequencing.

## Author contributions

ZW, SL, and BZ designed the research; ZW, JP, PB, and WS performed the experiments; ZW, KB, and SL analyzed the data; ZW, KB, QS, and SL wrote the paper with input from all the authors.

## Acknowledgments

The authors would like to acknowledge the support of USDA-NIFA Grant 2020-70410-32900 for the nanopore sequencing device. Kassaye Baley would like to acknowledge the support of GBCB program from Virginia Tech.

## Figure Legends

**Table S1** Structural variant distribution across 29 soybean genotypes, encompassing deletions (DEL), insertions (INS), inversions (INV), duplications (DUP), and inversion-duplications (INVDUP)

**Table S2** Structural variant length distributions across the 29 soybean genotypes

**Table S3** Structural variant distributions across different genomic regions across the 29 soybean genotypes

**Table S4** Statistical analysis of SV(s)-carrying genes associating with protein content, oleic acid content, methionine content, seed weight, plant height, and flower color using Benjamini-Hochberg test

**Table S5** Number of significant genes associating with selected agronomic and seed composition traits by Benjamini-Hochberg statistical test

**Table S6** The sequences of primers used in this study

## Methods and Materials

### Plant material

We chose 29 Glycine genotypes (Table S1) as the sequencing representatives for this study. All the plants were grown in the Kentland farm, Blacksburg, Virginia in the year of 2021.

### DNA isolation for Oxford Nanopore Technology (ONT) sequencing

High-molecular-weight DNA was isolated using DNA isolation protocol modified from Alonge et al ^1^. Five grams of young soybean leaves from 4-6-week-old plants were harvested and frozen by liquid nitrogen. Frozen leaf material was ground to fine powder using a mortar and pestle and transferred to a 50 mL Falcon tube. A total of 15 mL of pre-heated lysis buffer 1.4 M NaCl, 100 mM Tris pH 8.0, 2% CTAB (Hexadecyltrimethylammonium bromide, w/v), 20 mM EDTA, 0.5% Na_2_S_2_O_5_ (w/v), 2% 2-Mercaptoethanol (v/v, added freshly)) was added into the 50 mL Falcon tube. The lysate was incubated for 20 min at 60°C. A 15 ml chloroform/isoamyl alcohol (24:1) was added to the lysate to allow proper separation of the organic phase and aqueous phase and keep DNA protected into the aqueous phase. After centrifuge, a 12 cold isopropyl alcohol was added to precipitate the High-molecular-weight DNA. The eluted DNA was treated by RNase at 37 °C for 1 hour to degrade the RNA in the elution. The solution of chloroform/isoamyl alcohol (24:1) was then utilized to remove the RNase. After washing by 70% ethanol twice, the DNA was finally eluted by 1 X TE buffer (10 mM Tris, pH 8.0 and 1 mM EDTA). The gel-electrophoresis, qubit (Thermo Fisher), and tape-station (Agilent) were used to determine the concentration and quality of the DNA.

### Library preparation for ONT sequencing

Two ug of DNA was used to prepare the sequencing library, using the ligation sequencing kit SQK-LSK110 according to the manufacturer’s recommendations. Genomic DNA was subjected to end repair (New England Biolab Inc). After a bead clean-up (Applied Biosystems), sequencing adaptors were then ligated to the end-repaired DNA. Finally, the adaptor ligated DNA was once again subjected to bead cleaning. The DNA library was finally loaded onto an Oxford Nanopore PromethION flow cell for sequencing at Virginia Tech core facility, Genomic Sequencing Center.

### SV calling and sorting

For each of our 29 soybean genotypes selected for this study, ONT sequencing technology was used to generate long reads. Raw reads were basecalled on a GPU using Oxford Nanopore Technologies’ guppy base caller v. 5.0.11 with parameters --flow cell FLO-PRO002 --kit SQK-LSK110. Raw FAST5 files obtained from a single flow cell were then concatenated into a single file which was used for downstream analyses. Concatenated raw reads were aligned to the Wm82.a4 soybean reference genome ^3^. The Wm82.a4 soybean reference genome is a recently published preprint that improves to the previous Wm82.a2 soybean reference genome. This new reference genome is the most complete and accurate representation of the Wm82.a2 reference genome to date. The reference gene models used in this study, are the accompanying reference gene annotation set. Structural variants were called relative to this reference genome from aligned reads with MINIMAP2 (v2.23, --MD -Y -ax map-ont -t 50) and the resulting alignments were sorted and indexed using samtools (v 1.7, samtools view -bS, samtools sort - @ 20, samtools index -@ 20). To call SVs, we run Sniffles with parameters (sniffles -t 5 -s 20 -r 2000 -q 20 -d 1000 --genotype -l 30 -m, minimum read segment length for consideration = 1000, default = 2000). We chose relaxed parameters compared to the defaults because our samples are inbred cultivars and heterozygosity should therefore be nearly non-existent ^15^. As is convention, SV labels (insertions, deletions, duplications, inversions, translocations, and inversion-duplications) are defined with respect to this single reference genome and do not necessarily define the underlying mutations causing the genetic variation.

We applied a series of filters using bcftools ^24^ (bcftools view -i ‘(SVTYPE = “DUP” || SVTYPE = “INS” || SVTYPE = “DEL” || SVTYPE = “INV” || SVTYPE = “INVDUP” || SVTYPE = “TRA”) && ABS(SVLEN) > 49 && ABS(SVLEN) < 15000’) to remove any spurious calls that could affect downstream analyses. Any variants smaller than 50 nucleotides or larger than 15 kb were removed. We only retained deletions, insertions, inversions, duplications, and inversion-duplications for further analyses. We discarded unresolved breakpoints (SVTYPE=BND) as well as other complex types such as DEL/INV, DUP/INS, and INV/ INVDUP variants. We annotated all filtered structural variants based on their overlap with various gene features using R packages; GenomicFeatures, GenomicRanges and ChIPseeker ^25,26^. We retrieved the positions of the gene models for Williams82 assembly 4 from Phytozome and determined the location of peaks for each SVs in terms of genomic features of the following genic features: Promoter (<=1kb), Promoter (1-2kb), Promoter (2-3kb), Downstream (<= 300), and intergenic (3-15kb).

### Genomic validation

The selected structural variation candidates were validated by PCR using the corresponding genomic DNA as the template, where the WM82 genomic DNA served as the control. The PCRs were performed by *Taq* 2X Master Mix (New England Biolab Inc) according to the manufacturer’s instructions. Oligo primers are listed in Table S6.

### RNA isolation and real-time PCR

All RNA was extracted from soybean tissues using TRIzol reagent (Thermo Fisher Scientific) according to the manufacturer’s instructions and the method described by Wang et al ^20^. DNA residue was eliminated by treatment with UltraPure DNase I (Thermo Fisher Scientific). The integrity and quantity of total RNA were determined by electrophoresis in 1% agarose gel and a NanoDrop ND-1000 spectrophotometer (NanoDrop Technologies, Wilmington, DE). cDNA synthesis was performed using the SuperScript III First-Strand RT-PCR Kit (Thermo Fisher Scientific) with an oligo-dT primer based on the manufacturer’s instructions. Real-time PCR was conducted with cDNA as the template using the Quantitect SYBR Green PCR kit (Qiagen) according to the manufacturer’s protocol. Oligo primers are listed in Table S1. The soybean ELF1B gene was used as a reference gene, and data is presented as ΔCT.

### HPLC method to quantify kunitz trypsin inhibitor

The HPLC method for quantifying KTI was conducted in accordance with a previously established protocol ^27^. In brief, 10 mg of finely ground soybean seed powder was mixed with 1.5 mL of 0.1 M sodium acetate buffer (pH 4.5). Samples were vortexed and shaken for 1 h at room temperature. Following vigorous vortexing and a 1-hour incubation at room temperature, the sample underwent centrifugation at 12,000 rpm for 15 minutes. Subsequently, 1 mL of the supernatant was filtered through a syringe using an IC Millex-LG 13-mm mounted 0.2-mm low protein binding hydrophilic Millipore (polytetrafluoroethylene [PTFE]) membrane filter (Millipore Ireland).

The KTI in solution was then separated using an Agilent 1260 Infinity series (Agilent Technologies) equipped with a guard column (4.6 × 5 mm) packed with POROS R2 10-mm Self Pack Media and a Poros R2/H perfusion analytical column (2.1 × 100 mm, 10 μm). The mobile Phase A comprised 0.01% (v/v) trifluoroacetic acid in Milli-Q water, while mobile Phase B consisted of 0.085% (v/v) trifluoroacetic acid in acetonitrile. The injection volume was 10 μL, and the detection wavelength was set at 220 nm.

## Data Availability

Sequence reads generated in this study are deposited to NCBI Sequence Read Archive (SRA) database under BioProject accession number PRJNA918343.

